# Duoculture diminishes social interaction costs and improves growth rates of two fish species at different temperatures

**DOI:** 10.1101/2023.09.10.557086

**Authors:** Hong Yao, Paul. O’Bryant, Dean. Rapp, Zhe-Xia. Gao, Han-Ping Wang

## Abstract

Duoculture has been reported to increase growth rates of some fishes when reared in combination, due to “shading” effects between the species. Two experiments, one involving outdoor cage-rearing in a reservoir, and the other, indoor tank-rearing, were conducted within each of three temperatures ranges (means of ∼18.0°C, ∼22.0°C and ∼26.5°C), to determine whether duoculture of bluegill (BG) *Lepomis macrochirus* and yellow perch (YP) *Perca flavescens* would lead to improved growth relative to when the two species were reared separately. Juvenile bluegill and yellow perch were reared in triplicated groups each involving monoculture sets of 100% BG and 100% YP, and a duoculture set of 50% BG + 50% YP. Experiments in cages (Exp. 1) ran for 150 days while those in tanks ran for 126 days (Exp. 2). In Experiment 1, bluegill exhibited significantly greater (*P*<0.05) mean weight (*P*<0.05) in duoculture than in monoculture, under the high summer-like range of temperature (∼26.5°C) over most of the experiment, whereas yellow perch showed no significant difference in mean weight in duoculture versus monoculture. By the end of a 150-d experiment, bluegill in duoculture outweighed those in monoculture by 62.5%. In Experiment 2, yellow perch in duoculture grew significantly larger than in monoculture (*P*<0.05) under the warm thermal regime (mean of ∼22°C), while no significant differences were detected in mean weight of bluegill in monoculture versus duoculture. Yellow perch in duoculture outweighed those in monoculture by 33.1% at the end of the experiment. Yellow perch performed better in duoculture than in monoculture under the low thermal regime (mean of ∼18°C) in both experiments. A significantly greater reduction of CV_wt_ was observed for both bluegill and yellow perch in duoculture than in monoculture in Experiment 1, while no differences in CV_wt_ reduction were detected for bluegill in Experiment 2. Feed conversion ratios (FCR) of bluegill and yellow perch reared in duoculture were significantly lower than for both fishes reared in monoculture in Experiment 1, while there were no significant differences in FCR among the three groups throughout most of Experiment 2. Findings indicate that duoculture of yellow perch and bluegill holds good potential to improve growth and FCR, and to reduce size variation by diminishing social interaction costs.

## 1. Introduction

Competitive social interactions among fishes reared in groups have been shown to reduce food consumption, growth efficiency and ultimately, growth rates e.g., in Arctic char *Salvelinus alpinus* (1,2), bluegill *Lepomis macrochirus*, green sunfish *L. cyanellus*, and their hybrids ((3,4). Polyculture tends to impede interactions among conspecific fishes present in different rearing systems through “shading” which can be effective in reducing fish aggression and, in turn, social interaction costs, resulting in increased growth rates. Polyculture has long been practiced in many regions of the world to increase fish production. Many freshwater fishes and several marine fish species have performed better in polyculture and duoculture than in monoculture due to beneficial effects from the simultaneous presence of different species (synergism) and to changes in intra- or inter-species relationships and behavior (5–11). In a duoculture experiment with Atlantic salmon parr *Salmo salar* and two Arctic char strains, both species showed an increased growth rate relative to when each was reared in monoculture (12). In self-feeding systems, duoculture of rainbow trout and red-spotted masu salmon *Oncorhynchus masou macrostomus* reduced variability in self-feeder learning times (13).

Bluegill and yellow perch *Perca flavescens* are particularly important aquacultural and recreational species in the Midwest region and elsewhere in the U.S. Demand for these two high-value species has remained strong in the Midwest, and in the Great Lakes region particularly, for yellow perch. Recent concern over adequacy of commercial production has heightened the need to enhance growth rates of both species.

A recent study found that competitive social interactions substantially reduced food consumption, growth, and feed efficiency of hybrid bluegill (3). A follow-up study demonstrated that these costs were even higher for bluegill (4). The high social costs for bluegill may, to varying degrees according to culture setting, mask their higher growth potential (4). Yellow perch, like bluegill, is also an aggressive species. Anecdotal evidence indicates that yellow perch may establish dominance feeding hierarchies in ponds when feed is in short supply or offered to fish only in relatively restricted areas versus being widely broadcast.

Many aquaculture operations in the Midwest are small, family-run farms, which lack the capacity, either in land area or resources, to expand into larger-scale pond-culture operations. For many of these small farms, their continued existence will be possible only if pond cage-culture rearing is adopted. Development of methods to increase growth and survival of bluegill and yellow perch in cages will be essential to these farms. Duoculture of these two species shows promise as a method to reduce social interaction costs and enhance growth on a commercial scale.

The aim of the present study was to determine the efficacy of duoculture of bluegill and yellow perch in increasing growth rate and growth efficiency, by reducing social costs of both species under different temperature regimes and culture settings. Positive results from this research would have immediate impact on the economic return of many small aquaculture farm operations.

## 2. Materials and methods

### 2.1. Fish and experimental design

Two experiments were conducted in the Ohio Center for Aquaculture Research and Development at the Ohio State University.

*Experiment 1:* A cage-culture experiment was conducted in a reservoir during under spring and summer temperatures with large bluegill and yellow perch fingerling. Bluegill fry were secured from Ohio’s Hebron State Fish Hatchery, while yellow perch were self-propagated in indoor tanks and transferred to a nursery pond. The two species were cultured in distinct, 0.1-ha ponds for one growing season. When the fish reached similar sizes in the two ponds, they were harvested, whereupon they were randomly selected for experimentation. The experiment involved two phases: spring (Phase I) and summer (Phase II) and involved nine round cages (1.0m diameter) situated in the reservoir. The nine cages were positioned around a rectangular dock (4m × 8m) in the reservoir. An aerator was installed in the center of the dock and operated throughout the cage-culture rearing period. The experiment consisted of three fish groups, each with three replicate cages. The groups were yellow perch in monoculture (YPM), bluegill in monoculture (BGM) and yellow perch and bluegill together in duoculture (YPBGD). The stocking density was 250 fish per cage for all groups with a ratio of 50:50 for bluegill and yellow perch in duoculture. The mean weight of bluegill and yellow perch were 16.3 g and 19.1 g, respectively. All groups were acclimated for two weeks under experimental conditions before starting the experiment.

*Experiment 2*: This experiment was conducted in 55-L, indoor round tanks (0.5m diameter) under controlled temperature regimes for 18 weeks (126 days). Experimental temperatures were maintained at the same level (∼18°C) as in the first six weeks (Phase I) of Experiment 1 to evaluate the efficacy of communal rearing of the two species at low temperature in tanks by comparing with Experiment 1. Temperature was then increased to the warm thermal regime (∼22°C) and maintained at that level for 12 weeks (Phase II, Fig. 1). Both species were self-propagated in indoor tanks; bluegill were nursed in tanks while yellow perch were nursed in a pond. Yellow perch were feed-trained for three weeks before beginning the experiment. The experiment involved three groups as in Experiment 1, each with three replicate tanks. The stocking density was 70 fish per tank for all groups with a ratio of 50:50 for bluegill and yellow perch in duoculture. The mean initial weights of bluegill and yellow perch were 1.3g and 0.8g, respectively. All groups of fish were acclimated for two weeks under experimental conditions before the experiment began.

**Fig. 1.**
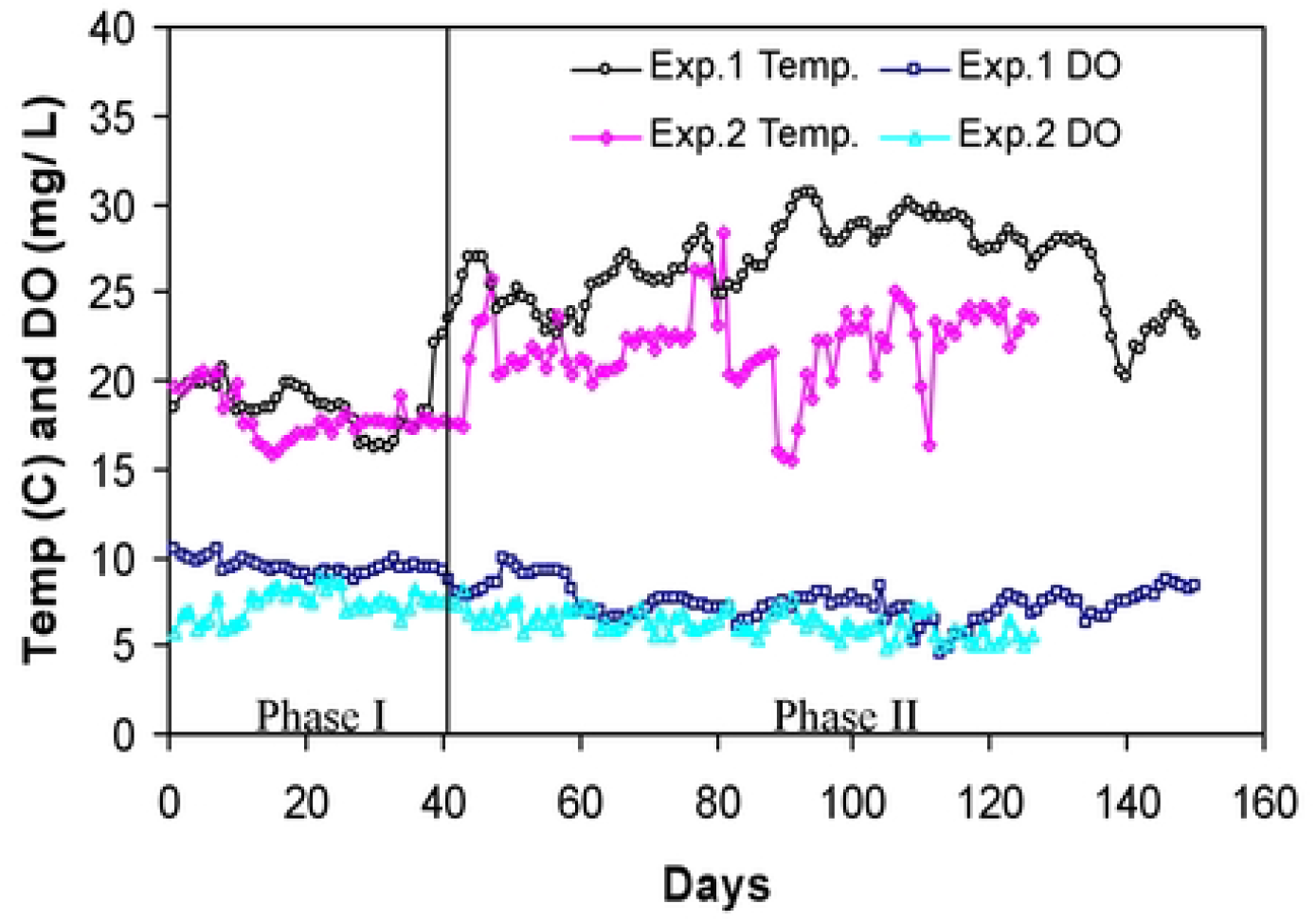
Daily vanances of the water temperature (Temp) and dissolved oxygen (DO) concentrations during Experiment I and Experiment II. The values for Experiment I are mean ofmoming and afternoon temperatures.

### 2.2. Feeding and water quality monitoring

In Experiment 1, fish were fed by hand to apparent satiation twice daily with a commercial trout feed (1.5 mm Silver Cup, 40 % protein, 11 % fat) at 09:00 and 16:00 hours, except during the extremely high temperature period (middle July to middle August) when fish were fed just once daily in the morning. During Experiment 2, fish were fed by hand to apparent satiation three times daily at 08:00, 12:00 and 16:00 hours using the same type of feed as in Experiment 1 but with a smaller pellet size. Dissolved oxygen (DO) and water temperature were measured twice daily in the morning and afternoon for Experiment 1 and once in the morning for Experiment 2, with a YSI 51B Dissolved Oxygen meter (Yellow Springs Instruments, Yellow Springs, Ohio). There were two distinct temperature phases in each of the two experiments. Mean water temperature was 18.4 ± 3.56°C (15.4 – 26.9°C) for the first 40 days, and 26.4 ± 2.47°C (20.0 – 30.6°C) for the remaining 110 days of the Experiment 1, and 18.0 ± 1.26°C (15.8 – 20.5) for the first 6 weeks and 21.9 ± 2.37°C (15.5 – 28.3) for the remaining 12 weeks of Experiment 2 (Fig. 1). The mean DO levels over the 150-d experiment were 7.3 ± 1.27 mg/L (4.8 – 9.7 mg/L) for the Experiment 1 and 6.6 ± 0.89 mg/L (5.1 – 8.8 mg/L) for Experiment 2 (Fig. 1). Feeding behavior, aggressiveness, and fish responses to scheduled handling procedures were examined closely.

### 2.3. Data collection and analysis

At the beginning and the end of the two experiments, fifty fish from each cage or tank were randomly sampled for weight and length. During the experiment, thirty fish were randomly sampled from each cage or tank for weight once every month for Experiment 1, and every three weeks for Experiment 2 to monitor growth. Food consumption by all fish in each cage or tank was determined weekly by weighing back remaining (if any) weekly feed allotments.

Absolute growth rate (AGR), feed conversion ratio (FCR) and coefficient of variation in weight (CV_wt_) were calculated over periods as follows:

AGR = (W_t_ – W_0_) × t^−1^, where t is the number of days of rearing, W_t_ is the mean body weight (g) at day t, and W_0_ is the mean initial body weight (g);

FCR = feed consumed per fish (g) × (mean weight gain (g)) ^−1^;

CV_wt_ = (100 × standard deviation) × (mean body weight) ^−1^.

Differences in mean responses in body weight, AGR and CV_wt_ among groups for the two fish species were tested by one-way analysis of variance (ANOVA) which was performed on ranked data and followed by Duncan’s test for group separation where appropriate.

## 3. Results

### 3.1. Growth in bluegill

In Experiment 1, no significant differences were detected in the mean AGR of cage-reared bluegill in duoculture versus monoculture during the initial, 40-d low-temperature period, although the mean weight of fish in duoculture was higher than those in monoculture on day 40 (Fig. 2A, B; Table 1). However, during the high-temperature summer period, mean weights of bluegill in duoculture were significantly higher (*P*<0.05) than in monoculture from day 41 to day 150 except for day 120 (Fig. 2A; Table 1). Significantly higher AGRs (*P*<0.05) were detected in duoculture than in monoculture during the period of D120 to D150 (Fig. 2B). By the end of the experiment, the bluegill in duoculture outweighed those in monoculture by 62.5%, with the growth rate in duoculture being nearly twice as high (1.87) as that for monoculture. A slightly negative growth in body weight in duoculture was observed at day 120 (2A, B).

**Table 1.** Mean ± SE of body weight at the initiation of the experiment and at its conclusion of Phase I and Phase II in the Experiment 1 and Experiment 2, along with coefficients of variation (CV) and differences in coefficients of variation (CV diff.), absolute growth rate (AGR). Means within a column in each experiment followed by different letters were significantly different (*P* < 0.05). Phase I: Day 0 – Day 40 for Experiment 1; Day 0 – Day 42 for Experiment 2; mean water temperature was about 18°C for both experiments. Phase II: Day 41 – Day 150 for Experiment 1 with a mean water temperature of 26.4°C; Day43 – Day 126 for Experiment 2 with a mean water temperature of 21.9°C.

|  | *Phase I* |  |  |  | *Phase II* |  |  |  |  |
| --- | --- | --- | --- | --- | --- | --- | --- | --- | --- |
|  | Mean BW<sub>0</sub><br>(g) | CV<br>(%) | Mean BW<sub>t</sub><br>(g) | AGR | Mean BW<sub>0</sub><br>(g) | Mean BW<sub>t</sub><br>(g) | CV<br>(%) | CV<br>diff. | AGR |
| Experiment 1 |  |  |  |  |  |  |  |  |  |
| BGD | 18.2 ± 1.67z | 61.74 | 31.0 ± 2.35z | 0.32 ± 0.07z | 31.0 ± 2.35z | 60.3 ± 3.40z | 44.47 | -17.3z | 0.24 ± 0.03z |
| BGM | 14.4 ± 1.05z | 69.11 | 22.4 ± 1.47yx | 0.20 ± 0.04zy | 22.4 ± 1.47yx | 37.1 ± 2.14y | 54.78 | -14.3y | 0.12 ± 0.02y |
| YPD | 19.0 ± 2.21z | 78.15 | 26.5 ± 1.84zy | 0.19 ± 0.09zy | 26.5 ± 1.84zy | 56.3 ± 2.81z | 33.56 | -44.6x | 0.25 ± 0.03z |
| YPM | 19.2 ± 1.57z | 77.26 | 21.3 ± 1.07x | 0.05 ± 0.05x | 21.3 ± 1.07x | 52.6 ± 2.13z | 38.36 | -38.9w | 0.26 ± 0.02z |
| Experiment 2 |  |  |  |  |  |  |  |  |  |
| BGD | 1.24 ± 0.09a | 80.3 | 3.01 ± 0.22a | 0.04 ± 0.005a | 3.01 ± 0.22a | 7.29 ± 0.70a | 80.7 | 0.4a | 0.04 ± 0.01a |
| BGM | 1.39 ± 0.06a | 57.7 | 3.30 ± 0.12a | 0.05 ± 0.003a | 3.30 ± 0.12a | 7.66 ± 0.30a | 55.9 | -1.8a | 0.05 ± 0.01a |
| YPD | 0.83 ± 0.04b | 46.6 | 3.91 ± 0.14a | 0.07 ± 0.004b | 3.91 ± 0.14a | 25.14 ± 0.86b | 34.7 | -11.9b | 0.25 ± 0.01b |
| YPM | 0.76 ± 0.28b | 53.4 | 3.31 ± 0.10b | 0.06 ± 0.002c | 3.31 ± 0.1b | 18.89 ± 0.47c | 36.0 | -17.4c | 0.19 ± 0.01c |

**Fig. 2.**
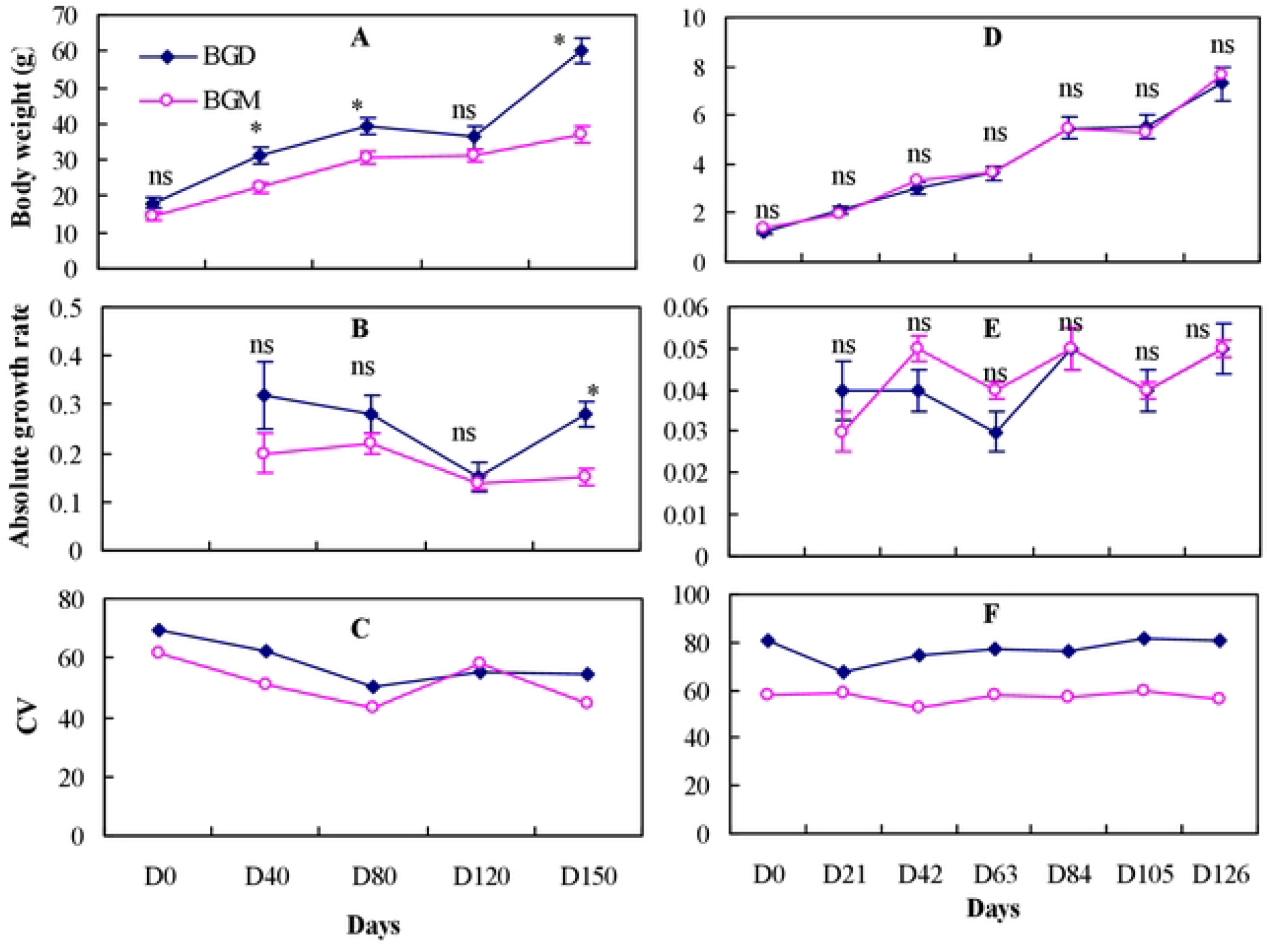
Standard body weight (g), absolute growth rate and coefficient of weight variation of the bluegill sunfish in Exp.I (left) and Exp. 2 (right) reared in monoculture (BGM) and duoculture (BGD). Values are means (±SE). Bars with the “ns” mean not significant difference and asterisk means significant difference *(P* < 0.05).

In Experiment 2, there were no significant differences in body weight or AGR of bluegill in monoculture versus duoculture during experimentation in tanks during the low- and warm-temperature periods.

### 3.2. Growth in yellow perch

In Experiment 1, significantly higher mean body weights and AGRs were detected for cage-reared yellow perch in duoculture settings versus in monoculture settings during the 40-day period of low temperatures (3A, Table 1). However, there were no significant differences in mean body weight and AGRs of cage-reared yellow perch reared in the distinct culture settings during 110-d period of high temperature (Fig. 3A, B; Table 1). Slightly negative growth in body weight was detected at day 120 in both duoculture and monoculture groups (Fig. 3A, B).

**Fig. 3.**
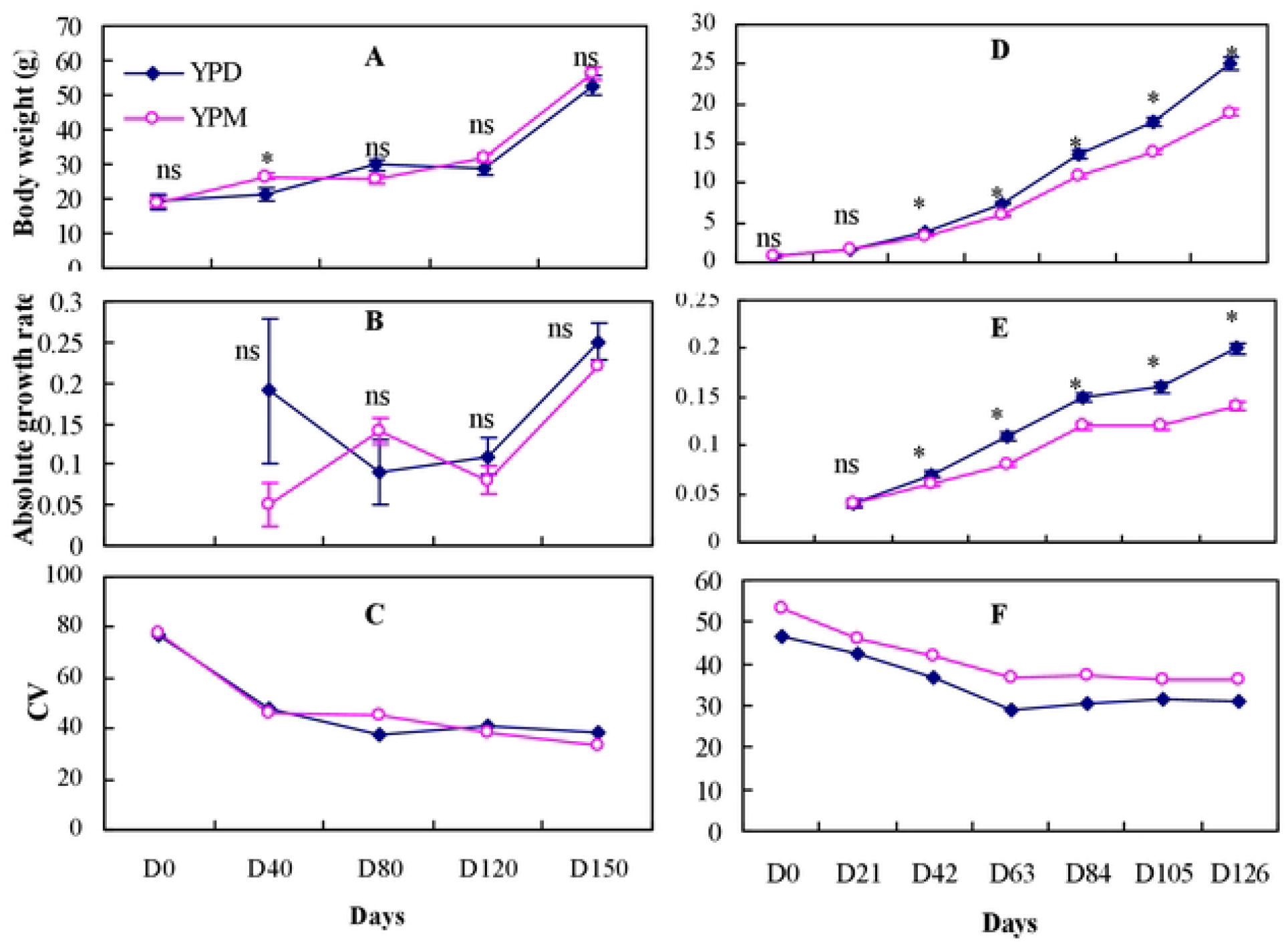
Standard body weight (g), absolute growth rate and coefficient of weight variation of the yellow perch in Exp.I (left) and Exp. 2 (right) reared in monoculture (YPM) and duoculture (YPD). Values are means (±SE). Bars with the “ns” mean not significant difference and asterisk means significant difference *(P* < 0.05).

In Experiment 2, mean weights of tank-reared yellow perch in duoculture were significantly greater than in monoculture system (*P*<0.05) in both Phase I and Phase II (Table 1), and over the entire 126-d experiment, except during the first 3 weeks (Fig. 2A); the yellow perch in duoculture outweighed fish in monoculture by 33.1% by the end of the experiment. Significant differences (*P*<0.05) were also detected in growth rates at the end of Phase I and Phase II (Table 1), and at the end of the entire experiment between the two types of culture systems (Fig. 2C, D).

### 3.3. CV changes

There were significantly greater reductions in CV_wt_ for both bluegill and yellow perch in duoculture than in monoculture in Experiment 1 (Table 1). In Experiment 2, a significantly greater CV_wt_ reduction for yellow perch was detected in monoculture than in duoculture, while there was no significant difference in CV_wt_ change in bluegill (Table 1). Overall, CV_wt_ for each species in both culture systems decreased slightly over the course of the experiments (Fig. 2C, 3C, F) except that CV_wt_ for bluegill was stable in Experiment 2.

### 3.4. Food consumption (FC) and FCR

At the end of Experiment 1, food consumption was significantly higher in the duoculture and bluegill monoculture groups than in the yellow perch monoculture group (Fig. 4A). The feed conversion ratio of fish in duoculture was significantly lower than in the bluegill monoculture and yellow perch monoculture groups (Fig. 4B). No significant difference was detected between the two groups of monoculture (Fig. 4B).

**Fig. 4.**
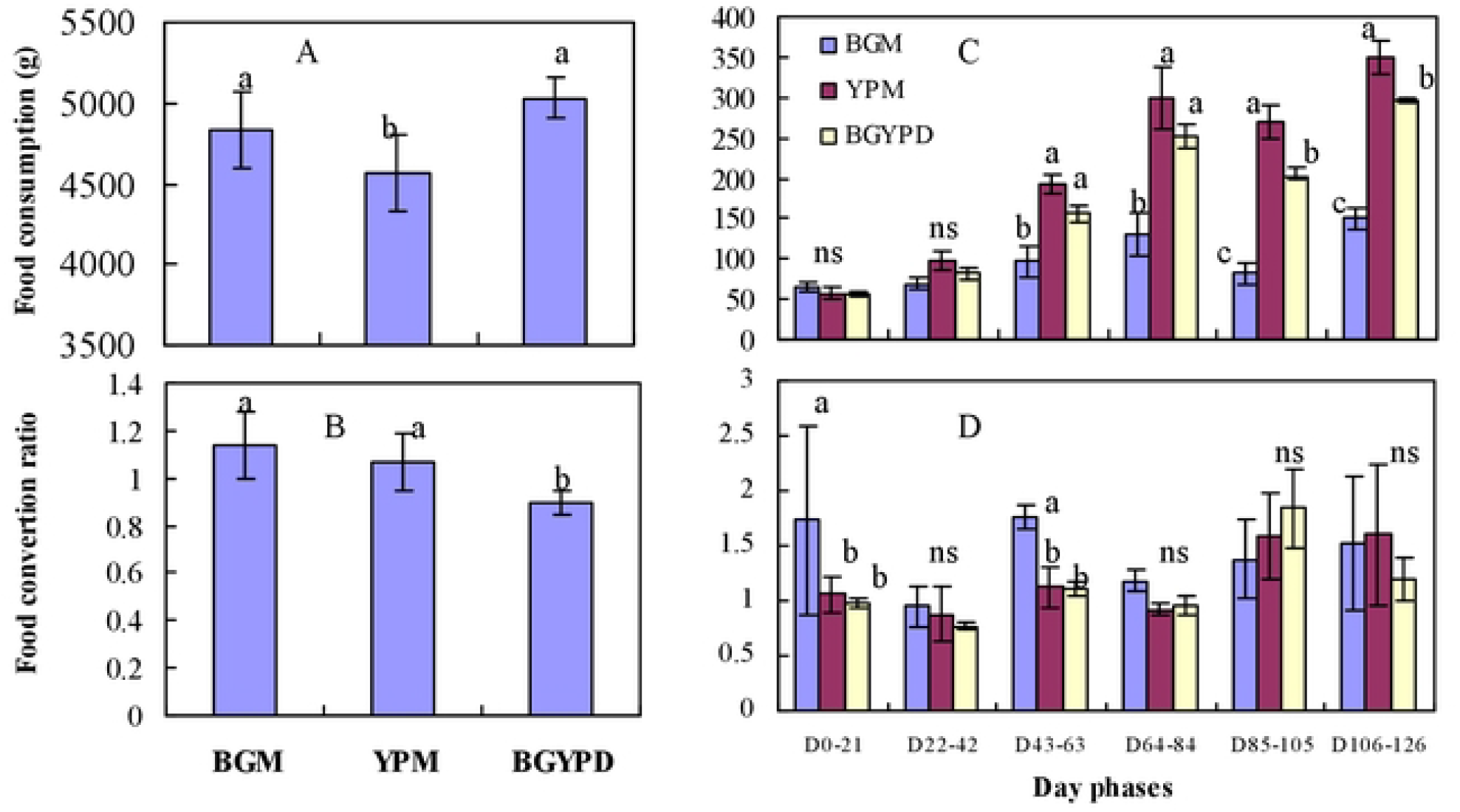
The food consumption and food conversion ratio (means± SE) of the bluegill and yellow perch in Exp. I (left) and Exp. 2 (right) reared in monoculture (BGM) and duoculture (BGYPD). Values with the different letters mean significant difference *(P* < 0.05) and ns means no significant difference *(P* > 0.05).

In Experiment 2, there were no significant differences detected in FC and FCR among the three groups in Phase I (the initial 6 weeks) with the exception of FCR during day 0 to day 21 (Fig. 4A, B). There were no significant differences in mean FCR among the three groups throughout most of Phase II. However, significantly more food consumption was detected in duoculture and in yellow perch monoculture groups than for bluegill in monoculture (4A, B).

### 3.5. Behavior

In monoculture, bluegill tended to remain in clusters near cage or tank bottoms. Aggressive interactions were largely intraspecific, occurring generally in corners and along cage or tank sides. In contrast, yellow perch dispersed homogenously the mid-level of the water column and showed less aggressive behavior than the bluegill. In duoculture, bluegill dispersed more randomly among the yellow perch. However, during the summer, particularly when temperatures were near 28 °C in July and August, yellow perch showed less movement and remained near the cage bottoms. Interspecific interactions were rarely observed.

## 4. Discussion

In the present study, bluegill and yellow perch reared together in duoculture each grew significantly larger than in the monoculture setting under certain conditions. Higher growth rates of fishes reared jointly in duoculture versus monoculture have been reported for many species combinations including common carp *Cyprinus carpio* with silver carp *Hypophthalmichthys molitrix* (14), Atlantic salmon with Arctic charr (12) and *Tilapia rendalli* with *Oreochromis shiranus* (7), and common carp with blue tilapia *O. aureus* (6,9). In contrast, duoculture of juvenile sharpsnout seabream *Diplodus puntazzo* and gilthead seabream *Sparus aurata* resulted in no growth improvement in recirculating aquaculture system (11) and growth loss in grow-out trials at a fish farm (10).

Bluegill and yellow perch are known to exhibit intraspecific aggression, especially juveniles, when the individual size variation is substantial in monoculture (3). As Holm (12) suggested, the mixing of species in duoculture could result in decreased intraspecific aggression via two avenues. In our experiment, duoculture of yellow perch and bluegill caused both species disperse more evenly, such that the mean distance between conspecifics was increased. Secondly, each species exerted a “shading” effect on the other, thereby reducing visual and physical contact which led to suppressed aggression. A “shading” effect has been suggested to underlie the improved growth that was observed in Atlantic salmon when reared in duoculture with Arctic char (5), and likewise in *D. puntazzo* when reared in duoculture with *S. aurata* (11).

Our findings indicate that the efficacy of duoculture of bluegill and yellow perch in improving growth rates is temperature dependent. Bluegill grew significantly larger in duoculture than in monoculture under summer thermal conditions (∼26.5°C), whereas yellow perch performed better in duoculture than monculture under the low temperature (∼18°C) and warm (∼22°C) temperature regimes. (15) found that the optimum temperature for bluegill growth was 30°C, and that growth rates of bluegill increased with temperature to approximately 30°C (16–18). The reported temperature range for yellow perch growth is approximately 11–25°C, with optimum temperatures being in the range of 22–25°C (19). Above 26°C, yellow perch exhibit signs of stress and reduced growth under conventional aquaculture conditions (19). During the high temperature period of Experiment 1 in summer, the mean water temperature was 26.4°C, which would have clearly favored growth of bluegill more so than yellow perch. Over the low-temperature periods of Experiments 1 and 2 (Phase I), the mean temperature was ∼18.0°C, especially during the last 12 weeks of Experiment 2, the mean temperature was ∼22°C. These temperature regimes were desirable for growth of yellow perch.

During the high-temperature period of Experiment 1, yellow perch in both monoculture and duoculture as well as bluegill in duoculture, showed slightly negative growth from July 15 to August 15, as detected on day 120. Two factors may account for this. First, the fish in all groups were fed only once a daily due to the high temperatures that occurred from day 90 to day 120. Secondly, high temperatures may have had negative effects on the growth and activity level of both yellow perch and bluegill. (19) indicated a lethal temperature of 32°C for yellow perch; at temperatures above 26°C they show signs of stress via behavior and reduced growth. Thus, disease and mortalities can be expected under conventional aquaculture conditions (19). Temperatures near 28°C appear sufficient to cause chronic stress in yellow perch as manifested by reduced feeding (20). Growth rates of bluegill have been shown to decline as temperatures increase above 30°C (16–18). Hence, we believe that the high temperatures experienced by yellow perch during the Experiment 1 may have reduced their activity and growth rates in all the groups. Admittedly, however, quantifying these effects is difficult. Our observations indicated that when experiencing temperatures close to 28°C, yellow perch tended to remain at the bottom of the cages and to reduce their activity. Moreover, we suspect that the inactivity of yellow perch in association with high temperatures, affected the growth of bluegill in duoculture. The lesser movement of yellow perch under high temperatures may have decreased their beneficial shading effect on bluegill which, in turn, increased bluegill’s intraspecific aggression and reduced their growth during the high temperature period. A sudden increase in CV_wt_ of bluegill in duoculture from day 90 to 120 provides further evidence of their increased social interaction costs in duoculture groups.

Both species showed declines in CV_wt_ in duoculture over the course of the experiment, whereas CV_wt_ has been reported to increase at times in polyculture (21). Significant CV_wt_ reductions for both bluegill and yellow perch in duoculture relative to monoculture in Experiment 1, indicated reduced social interaction costs in certain duoculture settings. However, a significant reduction of CV_wt_ for yellow perch was detected in monoculture but not in duoculture in Experiment 2. This may have occurred because some of the yellow perch in duoculture grew significantly faster than in monoculture, resulting in greater size variation. This result highlights the point that more may be occurring mechanistically than is reflected by simple quantification of CV_wt_ for improved growth rate in duoculture. More studies that focus on mechanisms that result in increased growth in duoculture settings are needed.

Food conversion ratio of fish in duoculture was significantly lower than that of bluegill monoculture and yellow perch monoculture in Experiment 1, indicating that social interaction costs are reduced in duoculture. The better FCR that was observed for yellow perch versus bluegill when the two monoculture treatments were compared, likely resulted from differences in social interaction costs between these two species. Moreover, the significantly lower food consumption of yellow perch in monoculture relative to bluegill in monoculture in Experiments 1 and 2, suggested that feed intake of bluegill contributed more to differentiations observed for significant feed consumption and utilization in duoculture population.

## 5. Conclusions

In conclusion, our study shows that bluegill and yellow perch each grew significantly larger in duoculture than in monoculture under different temperature conditions in both cage and tank settings. As market values of these two species are high in North America, especially in the Midwest, duoculture of yellow perch and bluegill shows promise as a method for improving growth and reducing social interaction costs at the commercial scale. Furthermore, the relative percentage of each species in relation to factors such as fish size, breeding time and production system, and optimal temperature conditions applied will need to be further investigated to optimize duoculture system for bluegill and yellow perch.

## Acknowledgements

This study was financially supported by the US Department of Agriculture. Salaries and research support were provided by state and federal funds appropriated to The Ohio State University. Thank Ohio Department of Natural Resource - Division of Wildlife’s Hebron State Fish Hatchery, and Mr. Ray Petering, Ohio Division of Wildlife’s Fisheries Supervisor, for providing the fish for this study.

